# Non-mechanical disruption of *Bacillus subtilis spores* allows sensitive and deep characterization of the minimal proteome for resuming a cellular lifestyle

**DOI:** 10.1101/2024.03.08.584050

**Authors:** Yixuan Huang, Alphonse de Koster, Zhiwei Tu, Xiaowei Gao, Winfried Roseboom, Stanley Brul, Peter Setlow, Gertjan Kramer

**Affiliations:** Laboratory for Mass Spectrometry of Biomolecules, Swammerdam Institute for Life Sciences, University of Amsterdam, Science Park 904, 1098 XH Amsterdam, The Netherlands; Molecular Biology & Microbial Food Safety, Swammerdam Institute for Life Sciences, University of Amsterdam, Science Park 904, 1098 XH Amsterdam, The Netherlands; Department of Molecular Biology and Biophysics, UConn Health, Farmington, CT 06030-3305 USA

**Keywords:** sample preparation by easy extraction and digestion, *Bacillus subtilis*, proteomics, spores, bacteria, single pot solid phase enhanced sample preparation, one pot, LCMS, timsTOF

## Abstract

In response to extreme conditions, *Bacillus subtilis* generates highly resilient spores characterized by a unique multilayered structure. This confers resistance against various chemicals and enzymes yet adding complexity to the analysis of the spore proteome. As the first step in bottom-up proteomics, sample preparation poses a significant challenge. We assessed how an optimized protocol for sample preparation by easy extraction and digestion (SPEED) performed compared to previously established methods “One-pot” (OP) and single-pot, solid phase-enhanced sample-preparation (SP3) for the proteomic analysis of *B. subtilis* cell and spore samples. We found that SPEED outperformed both OP and SP3 in terms of peptides and proteins identified, moreover SPEED highly reproducibly quantified over 1000 proteins in limited input samples as low as 1 OD_600_ of *B. subtilis* cells and spores. SPEED was applied to analyze spore samples of different purity by applying sequential purification following harvesting of spores. Comparison of the differential abundance of proteins revealed clusters likely partially stemming from remaining vegetative cells in less purified spore samples. We show that ranking of absolute protein abundance in cellular and spore samples further enables us to rationally differentiate integral spore proteins from vegetative remnants. This is of importance in applications and organisms where highly homogenous spore samples are difficult to obtain. A deep proteomic analysis of spore and vegetative cell samples with the new approach led to the identification of 2447 proteins, 2273 of which were further quantified and compared between *B. subtilis* spores and cells. Our findings indicate that pathways related to peptidoglycan biosynthesis, glycolysis, carbon metabolism, and biosynthesis of secondary metabolites are shared between cells and spores. This corroborates and extends earlier work stressing that despite marked differences in their physiological states, spores preserve vegetative cell (core) proteins, essential for revival under conditions conducive to growth.

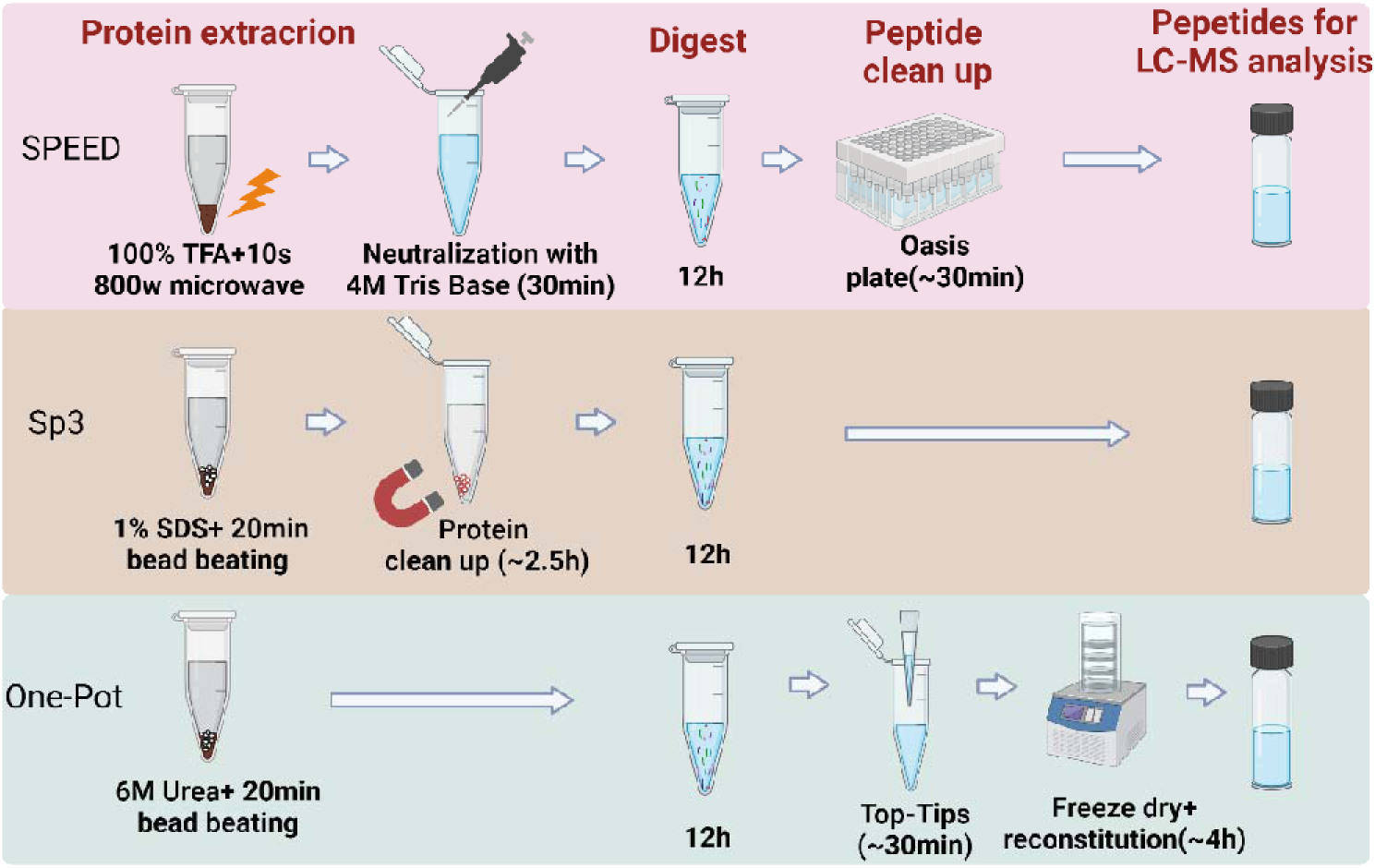

## Introduction

The endospores of *Bacillus* species are unparalleled in their resilience, upon encountering harsh environmental conditions, among all life forms on Earth. This resilience stems from a finely tuned interplay of genetic networks, culminating in the formation of these distinct stress survival structures. Sporulation is characterized by continuous protein turnover accompanied by numerous cellular adjustments (1). Distinct gene expression patterns control this process across the two compartments, mediated by the activation of alternative sigma factors, particularly σE, σF, σG, and σK. Initiated by starvation, the process of spore construction takes approximately 8 hours per cell (2), culminating in the creation of multi-layered spores resistant to threats such as heat, UV radiation, and various chemical agents. Upon detecting that the environmental conditions are suitable for growth, the spore can germinate and reactivate the vegetative cell cycle. The simplistic sporulation developmental program, genetic aptitude, and ease of genetic manipulation have made *Bacillus subtilis* a subject of considerable interest and the quintessential model for sporulation in bacteria. This is reinforced by its broad applications in biotechnology and agriculture, and its close relation to key pathogens such as *Staphylococcus aureus*, *Clostridioides difficile*, and *Listeria monocytogenes* (2, 3). in each developmental stage—be it cell growth, sporulation, or germination—proteins serve vital structural and physiological roles. With the increasing demand for in-depth knowledge of the molecular physiology of spore-forming bacteria, further elucidation of the biological processes at hand is needed to completely comprehend the metabolic and adaptive networks operating within *Bacillus* cells and spores. This is also why the global analysis of proteins by proteomics is increasingly used in the sphere of research of bacterial sporulation, germination and outgrowth (4–6). Even though *B. subtilis*, a well-studied model of gram-positive bacteria, was among the first to have its entire genome sequenced, only around 50% of its theoretical proteome is routinely measured in proteomics studies while up to 80% has been identified when aggregating data from a large number of studies (7, 8).

Spores, such as those formed by *B. subtilis,* are multi-layered structures that have evolved to survive threats to vegetative life by attributes such as a low-core water content and protective small molecules such as Ca-dipicolinic acid. Proteins are of particular importance as some, such as small acid soluble proteins (SASPS) and highly cross-linked coat proteins, play a crucial role in spore resilience and structure (9, 10). Because of this central role of proteins in the physical attributes of spores and concomitantly their resistance properties (11), analysis of protein content, quantity, crosslinking and complex formation and coat proteins using proteomics has been of the highest interest and multiple studies have been done over the past two decades (4–6, 12–16). Spore analysis through proteomics has multiple factors related to sample preparations that can influence the sensitivity, completeness and fidelity of the spore proteome: i) spore sample purity, ii) spore lysis efficiency, iii) protein extraction and solubilization efficiency and iv) completeness of protein digestion.

Spore sample purity directly influences the proteome detected as (significant) remnants of vegetative cells in the sample skew both the proteins detected as well as their quantity and will give a false impression of the spore proteome. Achieving high spore content samples by complete sporulation, killing of remaining vegetative cells and removal of cellular remnants through extensive washing and density gradient purification should be standard practice (17). However, achieving a completely pure spore sample can differ in sporulating bacteria as spores can have properties (size, tendency to aggregate) that make purification challenging (17). Spore lysis on the other hand directly influences sensitivity, as incomplete lysis will limit the amount of protein extractable from the spore sample. Given the unique structure of spores that have multiple rigid and highly crosslinked coat layers that give them increased resilience over the cell wall made by vegetative cells, lysis can be challenging. Spore lysis is achieved by rigorous mechanic disruption by zirconium aided bead milling to achieve comprehensive protein isolation (18). Complete extraction and solubilization of proteins following lysis is usually achieved using surfactants and chaotropic agents (e.g. SDS, SDC, Rapigest SF, ProteaseMax, Urea or Guanidine HCl). This ensures a stochastically accurate representation of all proteins of the spore proteome in the sample, ensuring that difficult to solubilize proteins (e.g. membrane bound and integral membrane proteins and coat proteins) are not underrepresented. This is particularly critical for *B. subtilis* spores, where the presence of an insoluble protein fraction (around 30%) poses a substantial challenge to comprehensive spore protein analysis (19).

A conventional procedure in a bottom-up proteomics experiment includes the extraction of complete protein sets, enzymatic digestion, chromatographic separation, MS detection and computational peptide identification and protein annotation (20). The bottom-up approach significantly depends on the complete breakdown of proteins into smaller peptides (21) as these serve as proxies for the protein amount in quantitative proteomics. As such, optimal conditions for digestion, along with all the factors mentioned above is a crucial last step before analysis in ensuring a balanced representation of the proteome as well as that an accurate quantification of proteins occur especially when aiming to get an overview of absolute abundance of proteins in a sample (22). Here strategies that help solubilize proteins like surfactants and chaotropic agents can also aid in tryptic digestion of difficult to digest proteins, even though high concentrations of denaturing agents will impair digestions by inactivating the proteolytic enzymes by denaturation, so a balance needs to be struck. Hence, comprehensive, and reproducible sample preparation methods are crucial, aiming at facilitating the generation of peptides from proteomes extracted from a diverse array of sample types.

To tackle the challenges posed by comprehensive spore proteomics we previously developed the “One-pot” method (OP), employing bead milling in 6 M urea under reducing conditions to disrupt intact spores of *B. subtilis*, *Bacillus cereus*, and *Peptoclostridium difficile* as well as vegetative cells of these organisms (19). Here the application of a chaotropic agent in conjunction with not clearing insoluble materials from the lysate prior to digestion, aids in the confident identification of over 1000 proteins. This approach (in essence an in-solution digestion approach) is also amenable to other lysis agents used for in solution digestion approaches such as guanidine hydrochloride (GnHCl), the bile salt sodium deoxycholate (SDC) or commercial mass spectrometry compatible surfactants like Rapigest SF and ProteaseMax. Other modern sample preparation techniques that utilize supplementary tools such as filters and beads, aimed at facilitating protein immobilization and/or purification like in-StageTip (iST) sample preparation using 18 discs (23) and filter assisted sample preparation (FASP) (24) with their ability to use harsher surfactants are also potentially useful. Indeed we previously optimized the use of single pot sample preparation for proteomics (25) (SP3) based on magnetic beads used in microbiology proteomics analysis (26–28) for spore proteomics in *B. subtilis* and *B. cereus* using the stronger surfactant SDS to improve solubilization of membrane bound proteins and less soluble proteins (5, 12). Although we used both OP and SP3 successfully to analyze spore proteomes and even performed the first true multi-omics study (29) of bacterial spores, a drawback of current approaches is the reliance on mechanical disruption techniques (bead milling) that require a relatively large sample input and are less amenable to state-of-the-art high throughput omics approaches using 96 or 384 well formats.

To address these shortcomings, and enable future studies where screening the spore proteome of many organism or mutants will become feasible we adapted the recently reported sample preparation by easy extraction and digestion (SPEED) (30) and optimized it for use with minimal amounts of bacterial spores. SPEED negates the need for mechanical sample disruption as shown by Doellinger *et al.,* as lysis-resistant sample types, can be lysed in pure trifluoroacetic acid (TFA) and proteins extracted efficiently by complete sample dissolution, while other interfering macromolecules such as DNA are hydrolyzed. Concomitantly with our study in spores, SPEED has been applied to show that it can be very useful in the high throughput analysis in a large panel of bacteria (30, 31). In the current study we compare the utility of SPEED to prepare proteomes from both *B. subtilis* cells and spores and compared these to our established OP and SP3 protocols regarding peptides and proteins detected, as well as surveying if there are any biases in protein classes quantified. We also show the utility of SPEED sample preparation for analyzing diminishing amounts of cells and spores towards higher throughput and single bacterial cell and spore proteomics. Finally with our optimized protocol we explore a longstanding question in spore proteomics, i.e. how do remnants of vegetative cells influence the apparent spore proteome. By analyzing spores of increasing purity alongside vegetative cells we show an easy way to distinguish between bona-fide spore proteins from those stemming from diminishing vegetative contamination. Moreover by correlating protein copy number from intensity based absolute quantitation (iBAQ) from label free quantitative proteomics in vegetative cells with proteins detected in common in spores, we show an additional way to provide a rational differentiation between likely vegetative cell contaminants from true spore proteins This is of particular interest when analyzing spores from organisms which yield higher amounts of vegetative contamination in spore samples and have spores that are difficult to purify by conventional spore purification approaches. Finally, we provide the most detailed quantitative and complete analysis of the minimal proteome spores need to resume a cellular lifestyle an classify the core groups of proteins present within these two stages of the bacterial life cycle.

## Materials and Methods

### Vegetative Cell Propagation and Sporulation

A single colony of *B. subtilis* PY79 was selected from an LB plate and grown in 3-5 ml LB medium (pH 7.5) at 37° C with a shaking speed of 200 rpm until the early log phase (OD_600_ 0.3-0.4) was achieved. Cell cultures were subsequently serially diluted in 5 ml of MOPS-buffered medium as previously described (32), and were subjected to overnight incubation at 37° C. One resultant culture exhibiting an OD_600_ between 0.3-0.4 was further diluted in pre-warmed MOPS medium and allowed to grow at 37° C in a 500 ml flask, subjected to a shaking speed of 200 rpm over a 3-day duration. Spores obtained through centrifugation were designated as “directly harvested spores (DH).” After centrifugation, these spores underwent three cycles of washing with chilled milliQ-water followed by centrifugation, thus resulting in what we termed “washed spores (WS)”. HistodenZ gradient centrifugation was employed to eliminate vegetative and phase-dark cells (33). Only samples that showed over 95% of phase-bright spores under microscopic observation were selected for further experiment as “ HistodenZ pure spore (HP).” In the case of vegetative cell preparations, cells were harvested from the LB medium when they reached the mid-exponential phase (∼OD_600_ 0.7). Three biological replicates were collected for spores and cells.

### Analysis of cellular and spore protein content and the relationship between cell and spore numbers and OD600 absorbance

The quantification of total protein concentration was performed using the Pierce BCA Protein Assay Kit from (Thermo Fisher Scientific, The Netherlands, Cat. No.: 23227). Following the manufacturer’s protocol, the BCA assay was implemented. Briefly, BCA standards were prepared with concentrations between 50 and 2000 µg/mL. 200 µL of working reagent (reagents A: B, 50:1) was combined with 10 µL of sample and standards, respectively. Absorbance at 562 nm was measured following a 30-minute incubation at 37°C. A standard curve was generated to calculate sample protein concentrations through a calibrated equation. Cell/Spore counting was undertaken by preparing 1 OD_600_ unit of cells and spores in 100 µL of phosphate-buffered saline (PBS) each. Following this, serial dilutions of 50-, 100-, and 200-fold were generated. A 10 µL aliquot of each diluted sample was loaded into a Bürker chamber, covered with a coverslip, and subjected to microscopic examination for cell counting after sedimentation was observed to be complete. Cell/Spore number was calculated by the following formula: total cells = total cells counted/100×4×dilution ratio×10^4^×0.1. Three technical replicates were performed for each sample.

### Sample preparation for mass spectrometry

#### Modified sample preparation by easy extraction and digestion (SPEED) protocol

Samples were prepared by a modified version of the SPEED protocol from *Doelinger et al* (30). Vegetative cells and spores were lysed in 30 μL 99% TFA (Biosolve; 10 min, room temperature) for each OD of *B. subtilis* sample and irradiated for 10 s at 800 W using a microwave oven at room temperature. Then samples were neutralized with 4 M Tris Base using 7 times the volume of TFA and further incubated at 70°C for 30 min after adding Tris(2-carboxyethyl) phosphine (TCEP) to a final concentration of 10 mM and 2-Chloroacetamide (CAA) to a final concentration of 40 mM. Subsequently, 100 µg (or the total amount of protein extracted for the experiment testing different starting amounts) of total protein were subjected to proteolysis with trypsin (protein ratio 1:50, 37 °C, overnight, with shaking at 400 rpm). Enzymatic digestion was stopped by addition of formic acid (FA) to a concentration of 2%.

Peptide samples were desalted using the Sep-Pak column containing 50 mg C18 cartridge (Waters, Milford, MA) or the 10 mg Oasis Plate (Waters, Milford, MA) depending on the estimated protein amount of the sample with a vacuum manifold. The desalting columns were conditioned with 1ml 100% acetonitrile (ACN) and 1mL of 60% methanol 1% FA in HPLC-grade water and equilibrated with 1% FA. Peptides were loaded on the column, washed with 1mL of 1% FA twice and peptides were eluted by the addition of 500 μL of 60% methanol 1% FA. The collected peptides were dried in a SpeedVac and stored at -80 °C until analysis.

#### Single pot sample preparation for proteomics (SP3) protocol

Spore/cell aliquots (∼400 μg) were re-suspended in 600 μL of lysis buffer (100mM Ambic, 1% SDS), and 400 μL volume of 0.1mm Zirconium beads was added (sample/beads 3:2 v/v). The tubes were put in a pre-cooled (4°C) bead mill and underwent ten 1-minute bead beating cycles of 30 seconds of milling at medium intensity followed by 30 seconds of cool down to limit thermal protein degradation. Afterwards, protein concentrations were quantified using the BCA assay per manufacturer’s instructions. Subsequently, TCEP and CAA were introduced to final concentrations of 10 mM and 30 mM, respectively, and the solution was incubated at 70°C for 30 minutes for reduction and alkylation. Proteins underwent purification using the SP3 protocol, after which trypsin was added at a protease-to-protein ratio of 1:50 (w/w), facilitating digestion at 37°C overnight. FA was introduced to achieve a final concentration of 1%, resulting in an approximate pH of 2. The sample was then subjected to centrifugation at 3000 x *g* at room temperature. The resultant supernatant was transferred to a new tube, centrifuged once more, and subsequently prepared for LC-MS analysis.

#### One Pot sample preparation protocol for bacterial cells and spores (OP) protocol

Samples were processed and digested following the OP method (19). Cells or spores were suspended in a 400 μL lysis buffer, composed of 6 M urea, 50 mM Ambic and 0.1mm Zirconium beads. Bead beating and BCA protein assay was done as described in SP3 protein clean-up. Subsequently, TCEP and CAA were introduced to final concentrations of 10 mM and 30 mM, respectively, and the solution was incubated at 70°C for 30 minutes for reduction and alkylation. The samples were subjected to enzymatic digestion with trypsin, employing a 1:50 ratio, and incubated overnight at 37°C. The digestion reaction was quenched with the addition of FA to pH < 4. Peptide containing supernatant was collected by centrifuging for 15 min at 15000 rpm. Peptide clean-up was done using C18 reversed-phase TT2 Top-Tips (Glygen) according to the manufacturer’s instruction.

### Basic reversed-phase (HpH) high-pressure liquid chromatography (HPLC) fractionation

The peptides from vegetative cell and spore samples were fractionated using a Waters CHS C18 1.9 μm particle size 2.1 × 100 mm column on an Ultimate 3000 HPLC (Dionex, Sunnyvale, CA, USA). The dried peptides were reconstituted in 100µl of 100 mM ammonium bicarbonate (Ambic) in HPLC-grade water. Peptides were loaded at a flow rate of 0.1 ml/min for 3 minutes and then eluted using a linear gradient (Solvent A: 10 mM Ambic, pH 10 in 5% Acetonitrile, 95% water, Solvent B: 10 mM Ambic, pH 10 in 90% acetonitrile, 10% water) from 3% to 40 % B in 34 minutes, then to 99% B in 2 minutes, kept at 99% B for 5 minutes before returning to initial conditions in 1 minute and re-equilibrated for 10 minutes. Peptide elution was monitored at 214 nm and the eluant was collected from 0-55 minutes into fractions using an automated fraction collector. Fractions were pooled into 10 pools, (i.e. fractions 1,11,21,31,41,51 pool1; 2,12,22,32,42,52 pool2; ….; 10,20,30,40,50 pool 10) freeze-dried and stored at −80 °C until analysis.

### LC-MS/MS workflow

The collected peptides were solubilized in a 35 μL solution of 0.1% formic acid (FA) and 3% acetonitrile (ACN). Subsequently, 5 μL of this solution was introduced into the Ultimate 3000 RSLCnano UHPLC system (Thermo Scientific, Germeringen, Germany). Peptide separation was performed on a 75 μm × 250 mm analytical column (C18, 1.6 μm particle size, Aurora, Ionopticks, Australia), which was maintained at a temperature of 50°C and operated at a flow rate of 400 nL/min with 3% solvent B for 3 minutes (solvent A: 0.1% FA, solvent B: 0.1% FA in ACN). Following this, a multi-stage gradient was applied, with 17% solvent B at 21 minutes, 25% solvent B at 29 minutes, 34% solvent B at 32 minutes, 99% solvent B at 33 minutes, kept at 99% solvent B till 40 minutes. The system was returned to initial conditions at 40.1 minutes and was held until 58 minutes for equilibration. The eluted peptides were electrosprayed by a captive spray source (Bruker, Bremen, Germany) via the column-associated emitter, and were analyzed by a TIMS-TOF Pro mass spectrometer. The instrument was operated in PASEF mode for standard proteomics acquisition, with quadrupole isolation widths set at 2 Th at 700 m/z and 3 Th at 800 m/z, and collision energy values varying between 20 and 59 eV across the TIMS scan range. Precursor ions possessing an m/z range from 100 to 1700 and a TIMS range from 0.6 to 1.6 Vs/cm^2 were chosen for fragmentation. PASEF MS/MS scans were initiated ten times with a total cycle time of 1.16 seconds, a target intensity of 2e4, an intensity threshold of 2.5e3, and a charge state range of 0-5. Active exclusion was enabled for a period of 0.4 minutes, with precursors being reevaluated if the ratio of the current intensity to the previous intensity exceeded 4.

### Data processing

The proteomics raw data was processed using Maxquant (version: 1.6.14.0) and matched with a UniProt proteome database (proteome ID UP000001570). Methionine oxidation was set as a variable modification, while cysteine carbamidomethyl was assigned as a fixed modification, allowing up to 2 missed cleavages by Trypsin. The “Match between runs” feature was selected with a matching time window of 0.2 minutes and the TIMS-DDA was chosen for group type. Classic normalization and a minimum ratio count of 2 were implemented for label-free quantification (LFQ), with all other parameters retained at default. Quantified protein data were reported in the proteinGroup.txt output file and were subsequently analyzed and visualized using Perseus (version 1.6.15.0). Data were filtered to remove potential contaminants, reverse hits, and protein groups “only identified by site”. Protein groups with more than 1 unique peptide were retained for further analysis. The predominant proteins were identified using the R/Bioconductor package *limma* with P value < 0.05 and absolute log2 fold change >1. Kyoto Encyclopedia of Genes and Genomes (KEGG) enrichment analysis of differentially expressed proteins was implemented using the ClusterProfiler R package (34).

## Results

### Comparison of methods using mechanical disruption (OP and SP3) versus chemical lysis (SPEED) for the analysis of cell and spore proteomes of *B. subtilis*

To ascertain how chemical lysis using SPEED performs in bacterial cell and spore proteomics against established methods using mechanical disruption we tested SPEED’s performance against that of OP and SP3 in *B. subtilis* cells and spores. Numbers of peptides identified by the different methods (**Figure. 1A**) show that SPEED sample preparation yields the largest number of identified peptides in both cells and spores (on average 13800 peptides in cells and 15993 in spores). Whereas the OP method identifies more peptides in spores compared to SP3, it identifies less peptides when applied in vegetative cells. In both cells and spores the peptides identified after SPEED sample preparation translated into the largest number of proteins, 1756 in cells and 1703 in spores (**Figure. 1 A, B**). Proteins identified in at least two replicates of one out of three method were further compared. 886 proteins in cells and 1095 proteins in spores were identified in all three methods (**Figure. 1C, D**). The OP approach distinctly identified 37 proteins in cells and 15 in spores, a significant contrast to the 298 cellular and 253 spore proteins identified solely by SPEED, and the 144 cellular and 82 spore proteins by SP3. The different unique proteins discerned by only a single methodology were assigned functional categories using *Subtiwiki* for analysis (**Figure. 1E, F**). In both cells and spores, SP3 extracts more membrane proteins than OP and SPEED. However, SPEED specifically identifies various other proteins related to coping with stress and regulation of gene expression.

**Figure 1.**
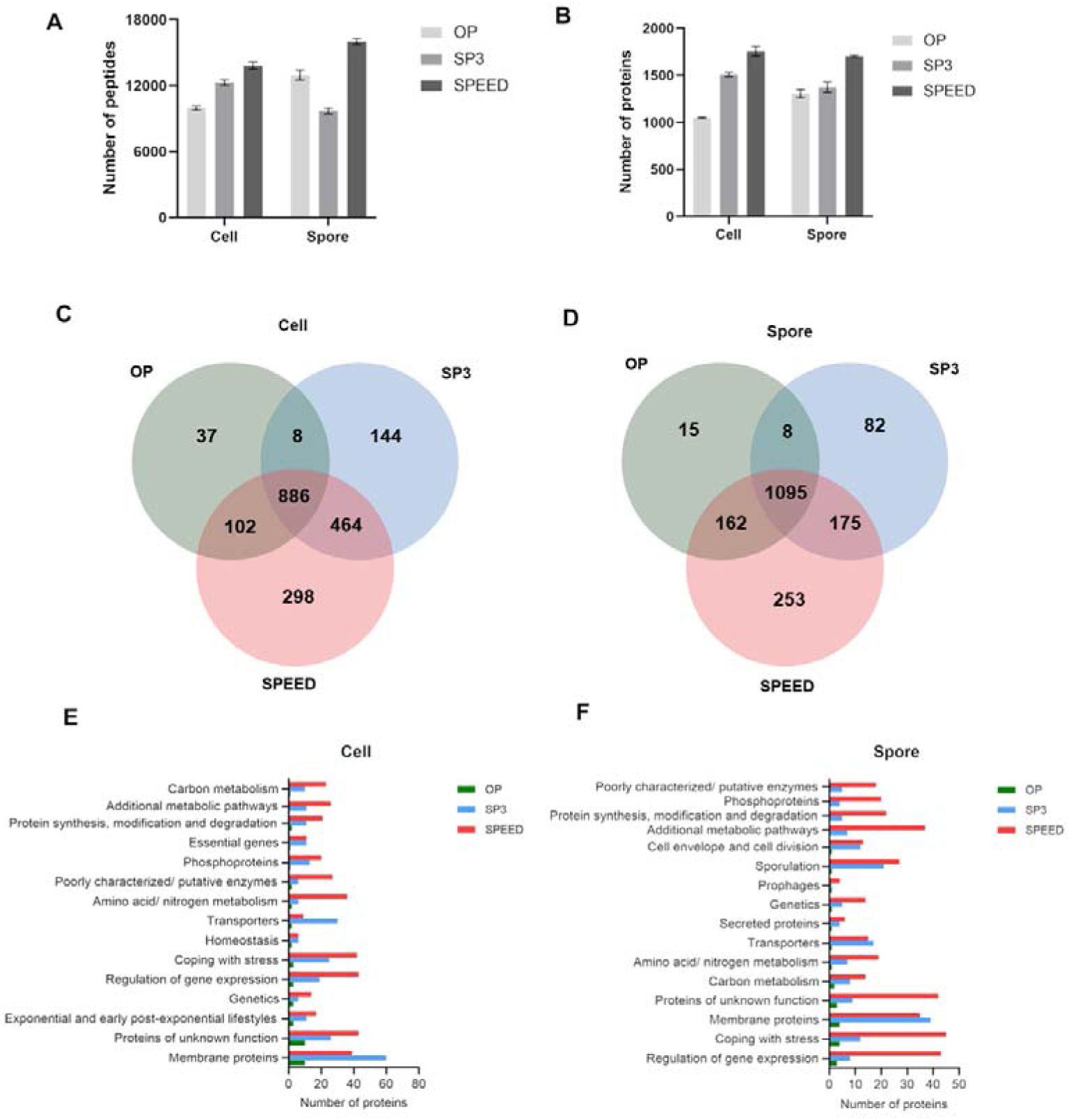
Comparison of OP, SP3, and SPEED regarding identified peptides and proteins. Number of peptides **(A)** & proteins **(B)** identified in *B. subtilis* cells and spores. Venn diagram of the identified proteins from the OP, SP3 and SPEED in *B. subtilis* cells **(C)** and spores **(D)**. (**E**, **F**) Functional analysis of proteins according to *Subtiwiki*. Three biological replicates were performed for each method.

To get an overview of the abundance levels of proteins quantified following sample preparation by the different methods, heatmaps were constructed using K-means cluster analysis (**Figure. 2 A, C**). In cells, SPEED extracted higher amounts of proteins in cluster K1 and K2, and lower amounts of proteins in cluster K3, compared to SP3 and OP. In spores, the SPEED extracted higher amounts of proteins in cluster K1 and K3, and lower amounts of proteins in cluster K2, than SP3 and OP. Furthermore, SP3 extracted higher amounts of proteins in cluster K2 than SPEED and OP. Using GOCC analysis on the different clusters (**Figure. 2B, D**), we found that the representation of membrane related proteins including membrane raft and integral components of the membrane is consistently higher in SP3 versus the other two methods in both cells and spores. Overall SPEED clearly outperforms the other two extraction and preparation methods regarding numbers of proteins detected.

**Figure 2.**
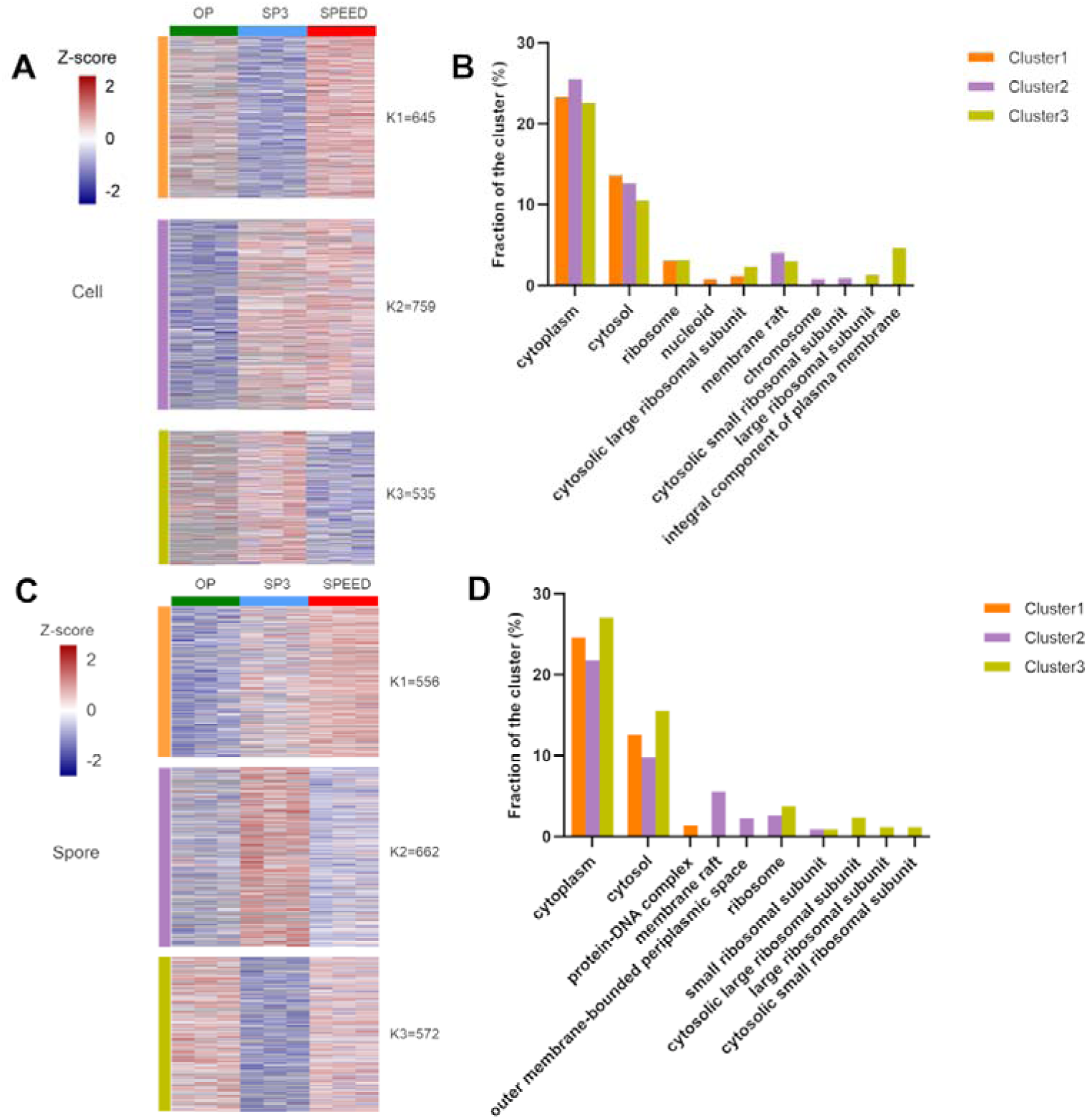
Heat map of identified proteins by OP, SP3, and SPEED and GOCC analysis of each cluster. Heat map of quantified proteins from cell samples **(A)** and spore samples **(C)**. Proteins were categorized into three clusters — K1, K2, and K3 — utilizing K-means clustering. Bar graph representing the GOCC functional analysis of clusters in cells **(B)** and spores **(D)**. Missing values in heat maps are indicated in grey.

To ascertain how SPEED performs using limited sample inputs we varied the OD_600_ of the starting material for SPEED sample preparation from 20 down to 0.1, correlating to numbers of vegetative cells ranging from 4.3x10^9^ and 0.2x10^8^, while it correlates to numbers of spores ranging from 2.0x10^9^ and 0.1x10^8^ (**Supplementary Table 1** and **Supplementary** Figures 1**, 2**). The coefficient of variation (CV) for quantitation of cellular proteins were similar, with median CVs of 0.077 (OD_600_=20), 0.066 (OD_600_=10), 0.064 (OD_600_=5), 0.069 (OD_600_=2.5) and 0.077 (OD_600_=1) (**Figure. 3A**). However, when the sample inputs were reduced to OD_600_=0.5 and 0.1, the median CVs rose to 0.114 and 0.194 respectively. Comparable results were observed with *B. subtilis* spores, with OD_600_=5 possessing the lowest median CV (0.084), while the median CVs for inputs of OD_600_=0.5 (0.116) and 0.1 (0.144) were higher compared to OD_600_=20 to 1 (**Figure. 3B**).

**Figure 3.**
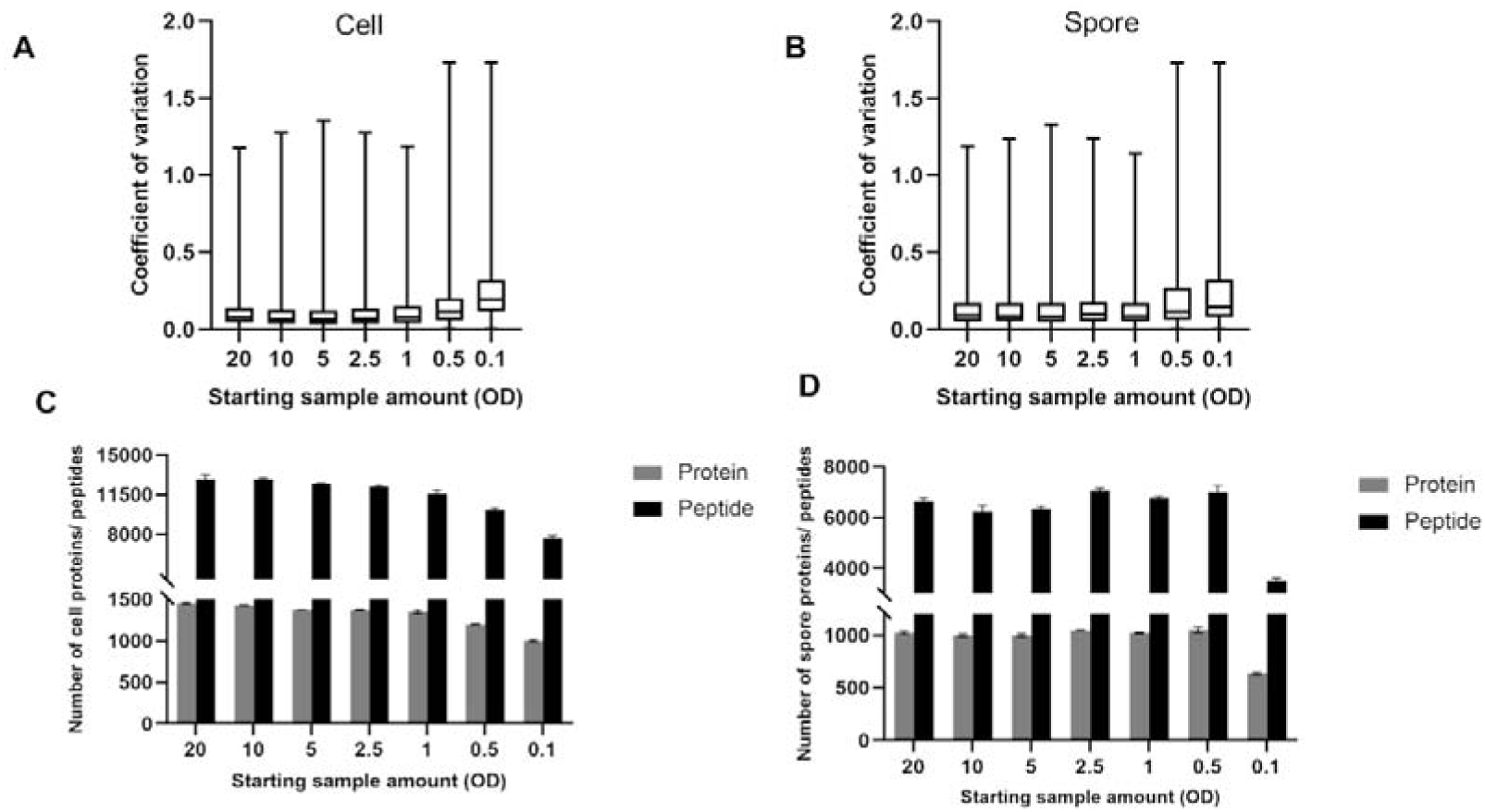
Comparison of results with different starting amounts of cells or spores using the SPEED extraction. Coefficient of variation (CV) of proteins and number of identified proteins and peptides in at least two replicates in *B. subtilis* vegetative cells **(A and C)** and pure HP spore samples **(B and D)**.

We further evaluated the number of extracted proteins and peptides. The number of cellular proteins identified was constant (ranging between 1461-1364) down to an input OD_600_ of 1 (**Figure. 3C, D**) going down notably when inputs of OD_600_=0.5 and OD_600_=0.1 were used (1204 and 1011 respectively). As the initial sample size decreased, the average number of detected peptides also decreased from 12877 to 7656. For spores, the lowest number of proteins was identified at OD_600_=0.1 with 697 proteins and 3502 peptides (average of 1041 proteins and 6678 peptides in other quantities of spores, **Figure. 3D**). Thus, for both *B. subtilis* cell and spore samples, OD_600_=1 seemed to be the minimal input required to achieve robust reproducible and adequate numbers of identifiable proteins. The numbers of identified proteins from 1 OD of cells and spores is similar to numbers reported previously for the OP method using much larger sample inputs combined with extensive fractionation (19). Overall, SPEED seems a promising method to enabling smaller amounts of bacterial sample for high-throughput applications for screening and time-series.

### The impact of spore harvesting procedures on the composition of proteome

When characterizing the spore proteome, a common complication is the heterogeneity of sporulation, which makes biological sense to the bacterium to hedge its chances for surviving adverse conditions. However, this means that spore samples are often a mix of mature spores and vegetative cells or vegetative (mother) cell remnants which result from the sporulation process. This complicates analysis as the resulting proteome contains both integral (encapsulated) spore proteins as well as proteins that adhere to the hydrophobic exterior of spores (from the mother cell or the environment of lysed cells) as well as remaining vegetative cells. However, to be able to discern bona-fide integral spore proteins from these other classes several purification approaches are often used in sequence. These purification approaches involve using the resistance and difference in physical properties of spores to kill and remove vegetative cells (through density gradient centrifugation) and washing steps to remove adherent proteins. To assess how different spore purification methods impact the final proteomic outcomes, we conducted a comprehensive comparison of three methods: direct harvest (DH), cold water washing (WS), and Histodenz purification (HP). The purity of the obtained spores varied across these methods, ranging from approximately 74.3% (DH) to 96.3% (HP) as shown in **Figure. 4 A** and **B**. Our findings revealed that the use of successive cold-water washing steps led to a slight improvement in spore purity, reaching 82.8%.

**Figure 4.**
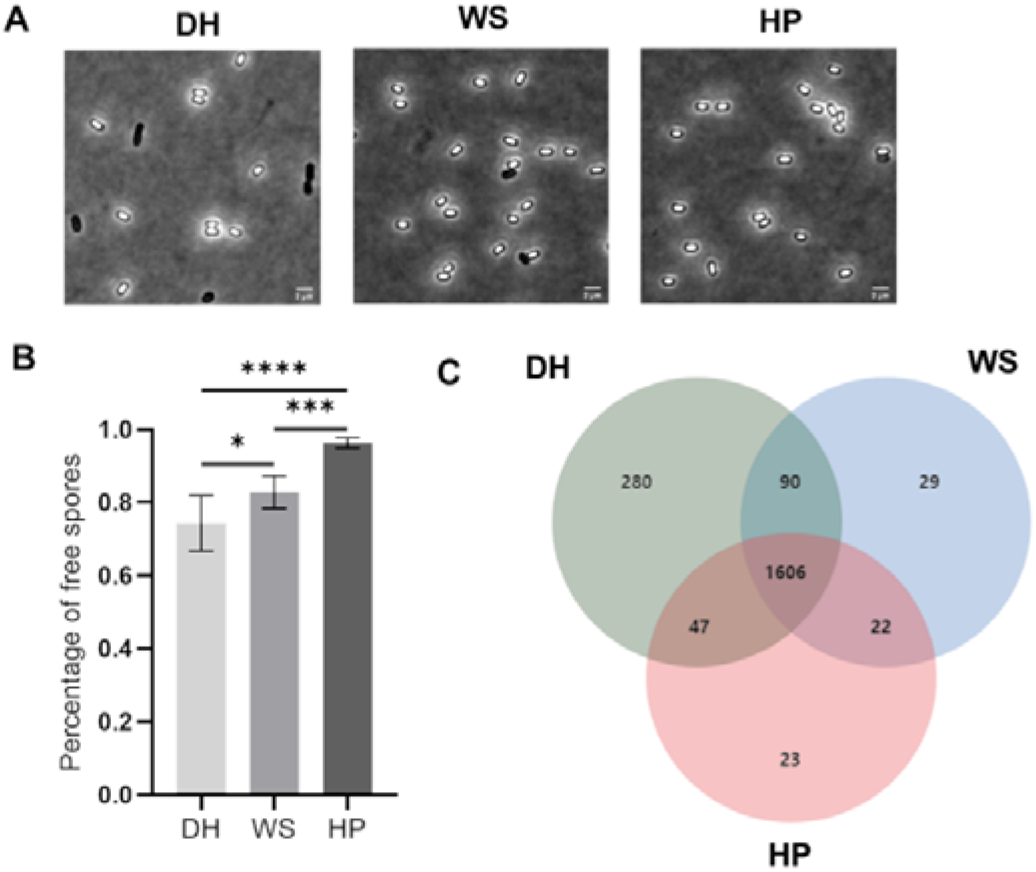
Comparison of different spore purification methods. Morphological images of spores purified with different methods **(A)** and the percentage of free spores **(B)**. **(C)** Venn diagram of identified proteins from spores purified differently.

These different purity samples were prepared through the SPEED protocol and analyzed by deep proteome analysis. Most proteins were identified in the DH sample (2023) followed by the WS (1747) and HP (1698) samples, which is expected, when background proteins stemming from unwanted sources are reduced (**Figure. 4C, Supplementary Tables 2**). Among all proteins identified, there were 1606 proteins identified in all three methods, while 280 proteins were only found in DH, 29 in WS, and 23 in HP. These sample specific proteins were classified according to *SubtiWiki* and are shown in **Table. 1**. With the DH method, there were 72 membrane proteins, which is the largest category, followed by coping with stress (54) and exponential and early post-exponential lifestyle (42). Additionally, 6 membrane proteins and 8 sporulation proteins were found in the WS and HP spores, respectively. All in line with obtaining a cleaner spore sample upon further purification.

**Table 1.** Classification of unique proteins from different spore purification methods according to *SubtiWiki*.

| Category | Number of proteins |  |  |
| --- | --- | --- | --- |
|  | DH | WS | HP |
| Membrane proteins | 73 | 6 | 4 |
| Coping with stress | 54 | 4 | 1 |
| Exponential and early post-exponential lifestyles | 42 | 0 | 0 |
| Proteins of unknown function | 37 | 2 | 3 |
| Regulation of gene expression | 37 | 6 | 4 |
| Additional metabolic pathways | 35 | 3 | 3 |
| Transporters | 28 | 3 | 0 |
| Cell envelope and cell division | 26 | 0 | 1 |
| Phosphoproteins | 24 | 2 | 1 |
| Protein synthesis, modification and degradation | 24 | 2 | 2 |
| Genetics | 24 | 0 | 1 |
| Secreted proteins | 18 | 1 | 0 |
| Carbon metabolism | 17 | 6 | 2 |
| Amino acid/ nitrogen metabolism | 16 | 5 | 1 |
| Sporulation | 16 | 4 | 8 |
| Nucleotide metabolism | 15 | 0 | 0 |
| Essential genes | 15 | 0 | 0 |
| Poorly characterized/ putative enzymes | 9 | 4 | 1 |
| Lipid metabolism | 9 | 2 | 1 |
| Homeostasis | 9 | 0 | 1 |
| RNA synthesis and degradation | 8 | 0 | 1 |
| Electron transport and ATP synthesis | 7 | 0 | 0 |
| Prophages | 4 | 1 | 1 |
| NcRNA | 1 | 0 | 0 |
| Detoxification reactions | 1 | 0 | 1 |
| Targets of second messengers | 1 | 0 | 0 |
| Universally conserved proteins | 1 | 0 | 0 |
| Efp-dependent proteins | 1 | 0 | 0 |

A deeper examination of the differences in protein abundance identified in the different purity samples was undertaken. Only those proteins identified in at least two replicates in at least one method were considered and quantified, which were categorized into different clusters based on K-means cluster analysis (**Figure. 5A**). In Cluster 1, most proteins are associated with sporulation, with markedly higher abundance in the HP sample compared to the other two (**Figure. 5B**), likely representing bona-fide integral spore proteins. Proteins extracted from DH were in high abundance in Cluster 2, consisting of 151 proteins encoded by essential genes, 145 protein synthesis, modification, and degradation, and 132 membrane proteins, notably the iBAQ rank of proteins derived from this cluster in vegetative cells is also notably higher (**Supplementary** Figure. 4). These proteins are reduced upon washing and Histodenz purification. Cluster 3 contained 137 membrane proteins, exhibiting higher intensities in both DH and WS only being reduced upon Histodenz purification. Our analyses also revealed a greater number of Spo0A-regulated proteins in clusters 2 and 3 compared to cluster 1 (**Supplementary** Figure. 3). This suggests that proteins derived from DH are not solely derived from spores but also from sporulating mother cells, a significant portion of which will likely be eliminated upon HistodenZ spore purification. Utilizing HistodenZ for spore harvesting effectively eliminates the majority of vegetative and sporulating cells. However, the resulting spores may not be entirely pure mature spores, as traces of mother cell proteins can still be detected, however comparing samples of differing purities can help differentiate between integral spore proteins (those showing increased abundance upon purification) and proteins that partially stem from different contaminating sources (those showing reduced abundance upon purification). Proteins which are reduced upon washing do not need to be solely contaminating proteins but may also both present in vegetative cells and spores. High abundance in vegetative cells (as shown by iBAQ rank in **Supplementary** Figure 4) combined with reduction during steps to eliminate vegetative cells is indicative of proteins measured likely stemming at least in part from vegetative “contamination” of the spore sample.

**Figure 5.**
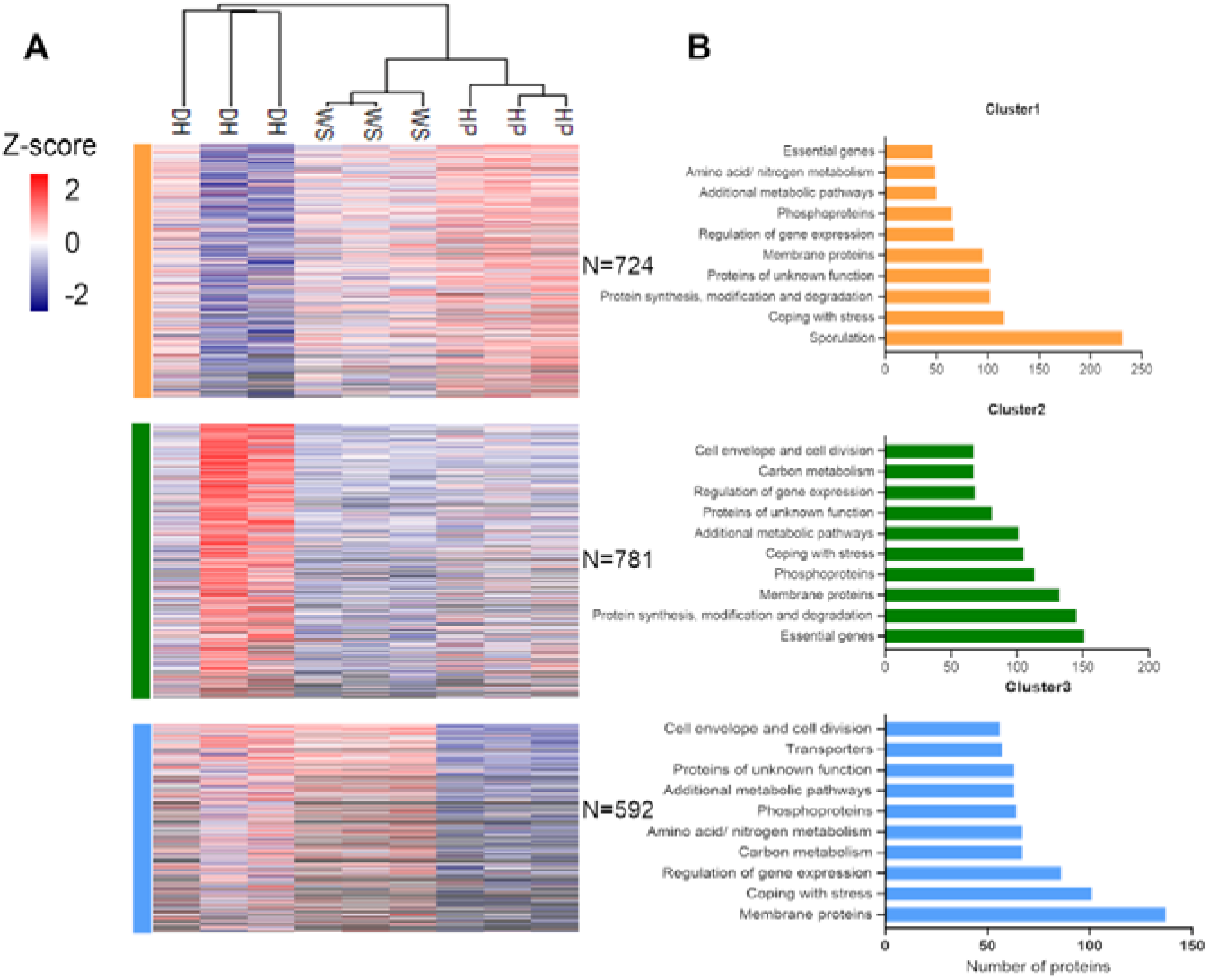
Deep proteome analysis of spores purified by different purification methods. Heatmap of the quantified proteins in at least two replicates in at least one method **(A)**. Proteins were assigned to 3 clusters with K-means analysis. **(B)** Bar plots of identified proteins from different clusters classified according to *Subtiwiki*.

### Comparative analysis of the proteome of vegetative cells and spores shows both the relative and absolute abundance of the minimal proteome to resume a cellular way of life

To facilitate an unbiased comparison of the cellular and spore proteome, we subjected equal protein amounts obtained by SPEED sample preparation to deep label free quantitative proteomics by peptide fractionation (10 fractions). The analysis of spore and vegetative cell samples yielded identification of 2330 proteins (**Supplementary Table 3**), among which 2273 were quantified across at least two of three biological replicates in *B. subtilis* spores or cells. Further comparison revealed 1439 proteins that were commonly identified in both spore and cell samples, representing 63.3% of the total proteins (**Figure. 6A**). Additionally, 575 proteins (25.3%) were found only in cellular samples, while 259 proteins (11.4%) were unique to the spores.

**Figure 6.**
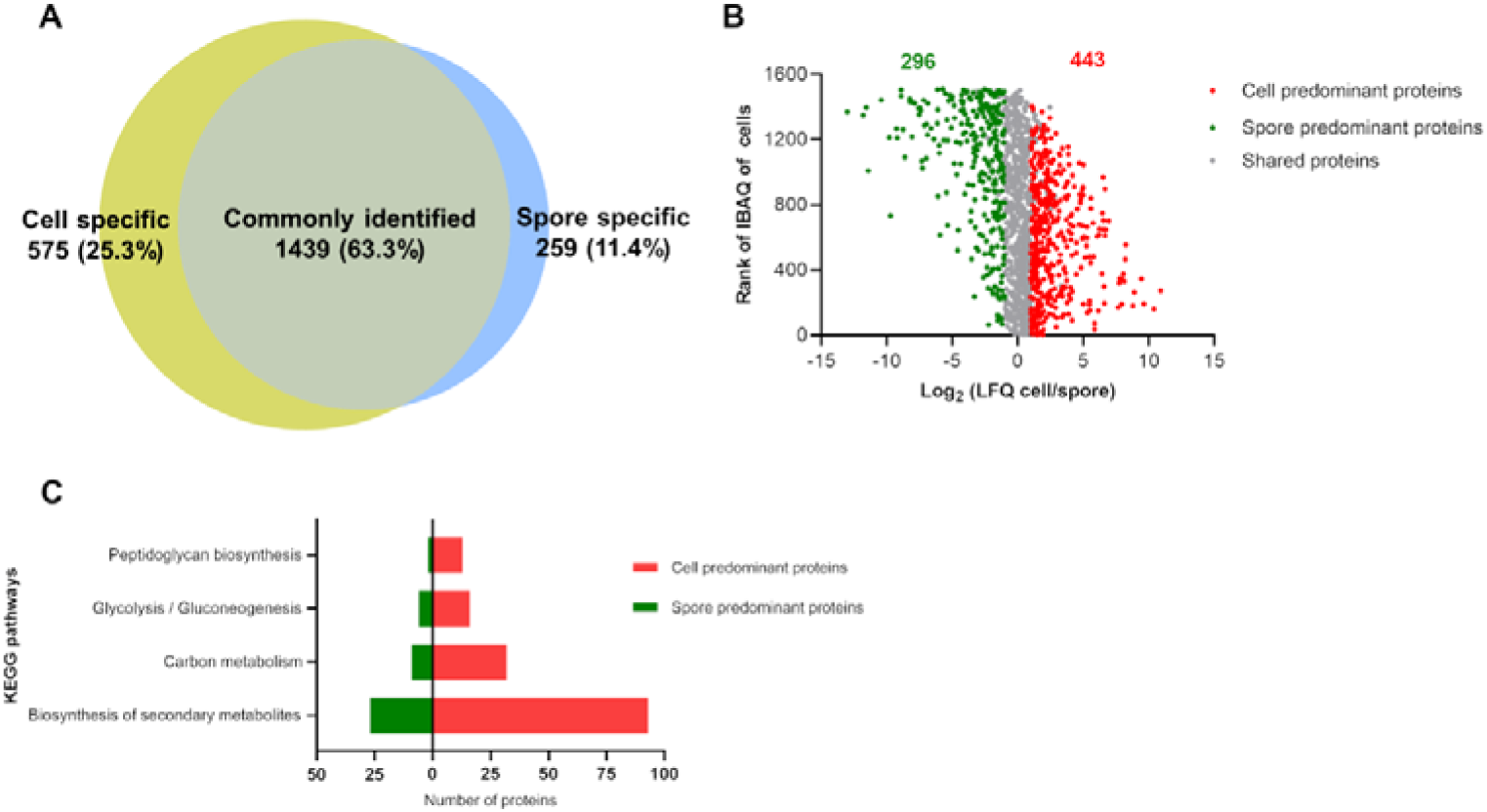
Deep proteome analysis of *B. subtilis* vegetative cells and HP spores. **(A)** Venn diagram of identified proteins from *B. subtilis* vegetative cells and HP spores. **(B)** Scatter plot of proteome comparison of *B. subtilis* vegetative cells and spores, y-axis shows iBAQ rank in vegetative cells with rank 1 denoting highest and rank 1600 denoting lowest protein copy number in cells. **(C)** KEGG pathway analysis of vegetative cell and spore predominant proteins; green and red bars represent the number of spore and cell predominant proteins, respectively.

Relative abundance of proteins in vegetative cells and spores versus absolute protein rank in vegetative cells are shown in figure 6B. The 1439 proteins that were quantified in both developmental stages were further analyzed to determine differential abundance between vegetative cells and spores. Proteins exhibiting a fold change > 2 and p value < 0.05 were considered differentially present between spores and cells. We found 296 proteins to be spore-predominant and 443 proteins to be predominantly present in vegetative cells. Remarkably, 700 proteins were shared between both spores and vegetative cells. To elucidate the roles of these predominant proteins in physiological processes, we conducted a KEGG pathway analysis. The most significantly enriched pathways were found to include biosynthesis of secondary metabolites, carbon metabolism, glycolysis/gluconeogenesis, and peptidoglycan biosynthesis (**Figure. 6C** and **Supplementary Table 4**). Further functional categorization of the shared, predominant, and specifically identified proteins was performed using DAVID. The proteins shared between spores and cells were mostly cytoplasmic proteins, transcription related proteins, and RNA-binding proteins and ribosomal proteins (**Table 2 and Supplementary Table 5**). Notably, spore-specific, and predominant proteins fell predominantly into the categories of sporulation, hydrolase, and oxidoreductase, whereas vegetative cell-specific and predominant proteins were mainly cytoplasmic and had transferase activity.

**Table 2.** Function of specific and predominant or shared proteins from *B. subtilis* cell and spore.

| UniProt keyword | Number of proteins |  |  |
| --- | --- | --- | --- |
|  | Cell specific and predominant | Spore specific and predominant | Shared |
| Sporulation | 0 | 107 | 0 |
| Cytoplasm | 223 | 0 | 173 |
| Capsid protein | 0 | 22 | 0 |
| Cell projection | 32 | 0 | 0 |
| RNA-binding | 0 | 0 | 64 |
| Virion | 0 | 22 | 0 |
| Cilium | 32 | 0 | 0 |
| Ribosomal protein | 0 | 0 | 41 |
| Hydrolase | 0 | 114 | 0 |
| Flagellum | 32 | 0 | 0 |
| Protein biosynthesis | 0 | 0 | 28 |
| Germination | 0 | 19 | 0 |
| Chemotaxis | 20 | 0 | 0 |
| Ribonucleoprotein | 0 | 0 | 41 |
| Protease | 0 | 34 | 0 |
| Bacterial flagellum biogenesis | 17 | 0 | 0 |
| rRNA-binding | 0 | 0 | 29 |
| Oxidoreductase | 0 | 65 | 0 |
| Bacterial flagellum | 13 | 0 | 0 |
| Glycolysis | 0 | 0 | 12 |
| Detoxification | 0 | 11 | 0 |
| Amino-acid biosynthesis | 51 | 0 | 0 |
| rRNA processing | 0 | 0 | 14 |
| Serine protease | 0 | 12 | 0 |
| Ligase | 49 | 0 | 0 |
| Ribosome biogenesis | 0 | 0 | 11 |
| Aromatic hydrocarbons catabolism | 0 | 9 | 0 |
| tRNA processing | 21 | 0 | 12 |
| Transcription | 0 | 0 | 77 |
| Lipid degradation | 0 | 7 | 0 |
| Cell division | 27 | 0 | 0 |
| Transcription regulation | 0 | 0 | 72 |
| Cell wall biogenesis/degradation | 42 | 23 | 0 |
| Methyltransferase | 33 | 0 | 0 |
| Isomerase | 0 | 0 | 33 |
| Cell cycle | 27 | 0 | 0 |
| Aminoacyl-tRNA synthetase | 0 | 0 | 12 |
| Peptide transport | 0 | 7 | 0 |
| Purine biosynthesis | 13 | 0 | 0 |
| Transcription termination | 0 | 0 | 4 |
| Secreted | 34 | 20 | 0 |
| DNA replication | 17 | 0 | 0 |
| DNA repair | 0 | 0 | 15 |
| Aminopeptidase | 0 | 6 | 0 |
| Pyrimidine biosynthesis | 10 | 0 | 0 |
| Metalloprotease | 0 | 10 | 0 |
| Transferase | 211 | 0 | 0 |
| DNA-directed RNA polymerase | 0 | 0 | 5 |
| Electron transport | 0 | 10 | 0 |
| Cell wall | 14 | 0 | 0 |
| Histidine biosynthesis | 0 | 5 | 0 |
| Primosome | 6 | 0 | 0 |
| Repressor | 0 | 0 | 41 |
| Peroxidase | 0 | 5 | 0 |
| Transducer | 9 | 0 | 0 |
| Nucleotide metabolism | 0 | 0 | 4 |
| DNA damage | 0 | 0 | 15 |
| Riboflavin biosynthesis | 0 | 0 | 5 |
| Flagellar rotation | 6 | 0 | 0 |
| mRNA processing | 0 | 0 | 3 |
| Cell shape | 23 | 0 | 0 |
| Helicase | 0 | 0 | 11 |
| Nuclease | 29 | 0 | 0 |
| tRNA-binding | 0 | 0 | 10 |
| Isoprene biosynthesis | 8 | 0 | 0 |
| Acetoin catabolism | 6 | 0 | 0 |
| Peptidoglycan synthesis | 18 | 0 | 0 |
| Nucleotide biosynthesis | 5 | 0 | 0 |
| Kinase | 57 | 0 | 0 |
| Teichoic acid biosynthesis | 7 | 0 | 0 |
| Allosteric enzyme | 9 | 0 | 0 |
| One-carbon metabolism | 6 | 0 | 0 |
| Two-component regulatory system | 29 | 0 | 0 |
| Gluconeogenesis | 5 | 0 | 0 |
| Restriction system | 5 | 0 | 0 |
| Lysine biosynthesis | 7 | 0 | 0 |
| Iron transport | 11 | 0 | 0 |

Previous studies into the spore proteome of *B. subtilis* were restricted to the analysis of relative abundance changes between spores formed at different environmental conditions or show the relative protein abundance between spores and vegetative cells as in the above. To gain insight into how different proteins play their respective roles in the dormancy and revival of these survival structures being able to ascertain absolute abundance of proteins globally rather than through targeted analyses in the dormant spores is of interest. Absolute abundance can add insight in both the stochastic composition of protein complexes and ascertain the abundance of stores of prepared proteins that might aid in the revival of the dormant spore upon receiving signals that conditions are favorable again for vegetative growth. Most proteomics studies look at relative protein level changes, as ascertaining a measure of absolute abundance of a protein is accompanied by several caveats. Nonetheless using intensity based absolute quantitation (iBAQ) or top 3 peptide quantitation (Hi3 peptide), several studies (22, 35–40) have shown that you can accurately estimate the absolute abundances of proteins using untargeted proteomics.

To show how various classes of proteins differ in their absolute protein abundance in spores and how this relates to their relative abundance between cells and spores, iBAQ versus relative protein abundance is shown in **Figure 7** and **Supplementary Table 6**. To show both shared, pre-dominant as well specific proteins we imputed a minimal value for proteins not detected in spores or cells and did not restrict the number of replicates a protein needed to be detected in to provide the broadest overview. This resulted in 1813 proteins which were categorized as cell predominant (460), shared (793) or spore predominant (298) using the same categorization as in the above. In addition, a category of spore specific proteins (262) and cell specific proteins (503) was added that had a fold change > 128 and p value < 0.05 (cell specific proteins are not shown in **Figure 7**). Protein abundance in spores was subsequently divided into 4 groups, i.e. high abundant (log10 iBAQ > 6.0 or the proteins above the 90^th^ percentile), abundant (log 10 iBAQ between 5.25 and 6.0 or between the 3^rd^ quartile and 90^th^ percentile), mid-range (log 10 iBAQ between 3.78 and 5.25 or between the 1^st^ and 3^rd^ quartile) and low abundant (log 10 iBAQ <3.78 or below the 1^st^ quartile). Overall, there is a trend that cell predominant proteins tend to have lower absolute abundance and spore predominant proteins tend to have a higher absolute abundance in spores (**Figure 7A**). However, **Figure 7A** also shows that there are many exceptions to this trend, and this underscores the need for absolute quantitation of protein levels as relative quantitation alone does not provide this same insight.

**Figure 7.**
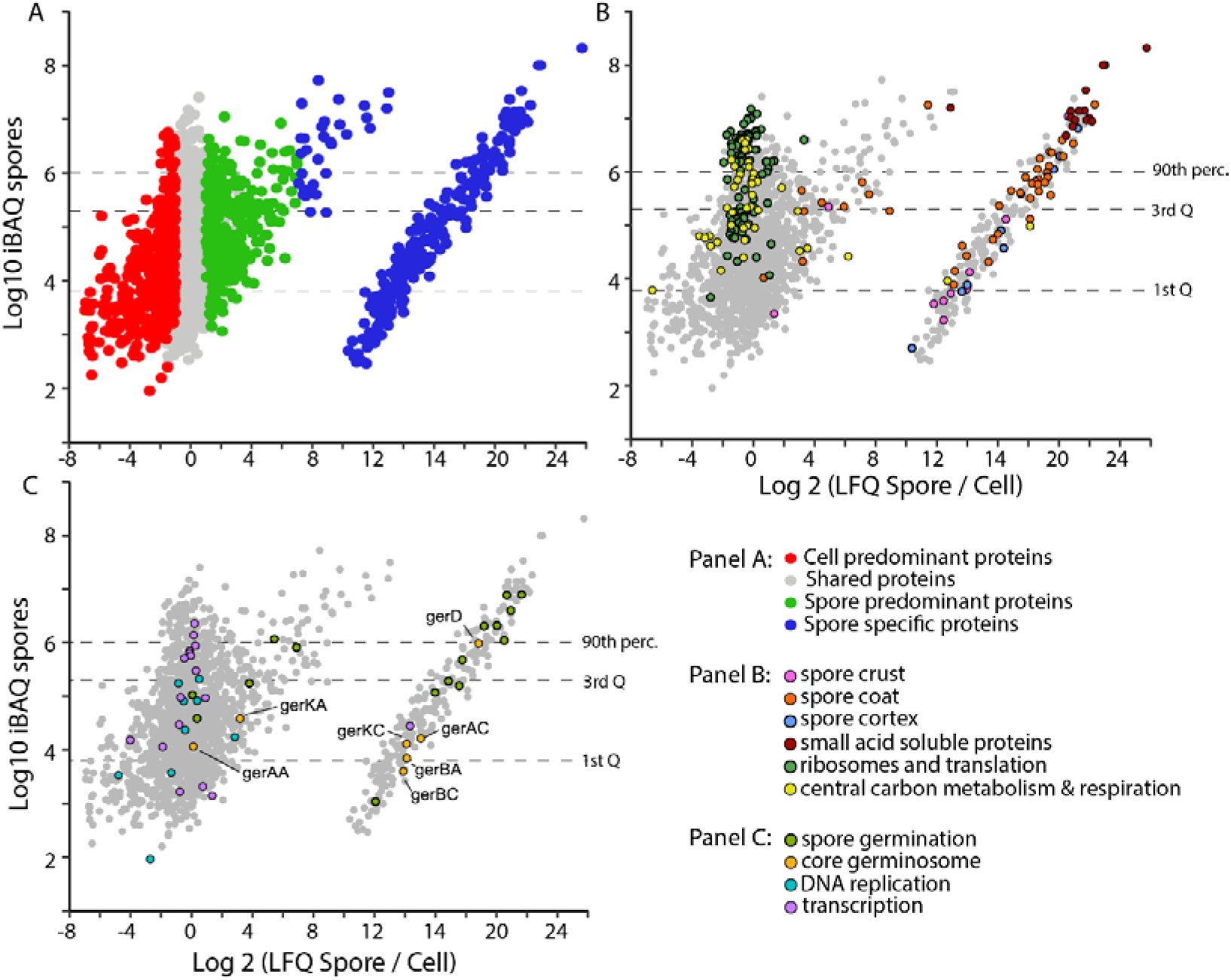
Profiling the abundance of proteins in spores shows the minimal proteome for resuming vegetative growth. Relative abundance (LFQ) of proteins in spores versus cells are shown on the x-axis, while corresponding absolute abundance of the proteins in spores is shown on the y-axis for all panels (iBAQ). Dashed lines show the 90^th^ percentile (90^th^ perc.), 3^rd^ quartile (3^rd^ Q) and 1^st^ quartile (1^st^ Q) of protein abundance in spores. These represent groupings of absolute protein abundance in the bacterial spore grouped in: highly abundant (>90^th^ perc.), abundant (90^th^ perc.<x<3^rd^ Q), mid-range (3^rd^ Q <x< 1^st^ Q) and low abundant (< 1^st^ Q) and can be found in **Supplementary Table 6**. (**A)** the relative abundance of proteins in spores versus vegetative cells are subdivided in cell predominant (log 2 fold change <-1, p <0.05), shared (log2 fold change -1≤x≤1,) and spore predominant (log 2 fold change >1, p<0.05) as described in figure 6, additionally a 4^th^ category of spore specific proteins (fold change >7, p<0.05) is added for proteins that are almost exclusively detected in spores (added by imputing minimal values for LFQ for proteins not-detected in vegetative cell samples the counterpart cell specific proteins are not shown but can be found in **Supplementary Table 6**. (**B)** proteins residing in specific spore layers are shown (see legend), as well as proteins needed to restart metabolism and protein synthesis upon germination which along with small acid soluble proteins are the most abundant proteins present in spores (residing mostly in the high abundant and abundant range of absolute protein abundance). **(C)** proteins involved in spore germination as well as the core germinosome have a varied abundance within the dormant spore, while proteins of core pathways such as DNA replication and transcription over all four abundance ranges.

Part of the spore’s resistance is thought to come from various types of proteins that associate with DNA in the core and are part of the cortex, coat and crust that form multiple protective layers around this spore core. Some of the most abundant proteins are the small acid soluble proteins (SASPs, shown in burgundy in **Figure 7B**) which as a class fall in the highly abundant part of the spore’s proteome. The proteins of the spore coat (orange in **Figure 7B**) are more varied in their abundance, while proteins localized in the crust and cortex (shown in pink and blue respectively in **Figure 7B**) mostly are mid-range to low abundant. Apart from protection and stability under harsh conditions the spore also contains proteins that enable it to germinate once conditions are favorable again. These so-called germination proteins (green in **Figure 7C**) vary in abundance from high to mid-range, while germinant receptors that make up the core Germinosome (light orange in **Figure 7C**) that is involved in nutrient sensing in germination are mostly low to mid-range apart from GerD, which is abundant in spores.

Strikingly proteins of the central carbon metabolism and respiration (yellow in **Figure 7B**), and the ribosomes and translation (green in **Figure 7B**) are also for a large part high abundant and abundant in spores on a par with many of the classically spore associated proteins of the core, cortex, coat, and crust. Proteins that facilitate transcription (purple in **Figure 7C**) are also high to mid abundant while DNA replication (aquamarine in **Figure 7C)** proteins range from mid to low abundance. All in all, many of the core pathways represented in vegetative cells also make up a significant part of the proteome of a dormant spore, whether they are there through active recruitment or through passive engulfment remains to be determined.

## Discussion

The past decade has seen significant advancements in tandem mass spectrometry, enabling a more profound characterization of microorganisms (41, 42). The wide adoption of bottom-up proteomics is largely attributed to the fact that peptides are more readily separable and identifiable by mass spectrometry than intact proteins (43). Especially with the use of label-free over labelled quantitative proteomics, efficiency and reproducibility are at the forefront of concerns when establishing a robust and sensitive proteomic workflow. Spore-forming bacteria, unlike other species, have a unique life cycle in which they form bacterial spores. These are characterized by their multilayer proteinaceous coats and thick peptidoglycan layer, which offers significant resistance to cell lysis and complete protein extraction (44). Thus, complete lysis and protein extraction require additional attention, when designing a proteomics workflow for spores. Although, the novel approach to sample preparation without mechanical lysis SPEED has been applied to various of biological materials amongst which are bacterial vegetative cells (31, 45, 46), its performance in bacterial spores has not yet been assessed. Comparative analysis of the protein profiles from *B. subtilis* cells and spores using SPEED sample preparation versus methods we established previously for bacterial spores (SP3 and OP) were first performed. This demonstrated the superior protein extraction by SPEED, compared to OP and SP3, which can be attributed to the enhanced lysis efficiency of pure TFA. However, when looking at differences in representation of protein classes SP3 exhibited superior efficacy in content of membrane proteins compared to both OP and SPEED. The use of SDS in SP3, a strong anionic detergent known for protein-denaturing properties, still enables better solubilization and denaturation of membrane proteins by breaking down hydrophobic interactions and solubilizing hydrophobic protein domains (47). This is an important aspect to take into consideration when choosing a sample preparation technique where the main objective is to detect membrane bound proteins, despite the overall superior performance of the SPEED extraction approach.

Developing a comprehensive bacterial sample preparation method without mechanical disruption is highly appealing, as it opens the way to high sensitivity (single cell) and high throughput proteomics using 96 and 384 well formats. This is of particular interest when targeting specific rare (sub)populations like persisters (48), bacteria isolated from the skin or gut microbiome (26, 49), and special bacterial samples or mutant libraries that can then be cultured and processed prior to analysis directly in 96 or 384-well plate formats(50). Such samples often present challenges for MS-based proteomics due to the difficulty in obtaining sufficient material for high-quality mass spectrometric data. SPEED is highly attractive in this sense, as we determined that with only 1 OD_600_ of *B. subtilis* cells or spores over 1000 proteins are identified with 80% of proteins having high quantitative reproducibility (CVs < 0.2) in both *B. subtilis* cells and spores. This likely stems from the non-mechanical disruption by TFA of cells and spores, which allows for the efficient extraction in small volumes in a single vessel of proteins without the losses associated with mechanical disruption and challenges in MS compatibility with the use of detergents and chaotropic agents (51).

In addition to being used as probiotics and antigen delivery vectors, *B. subtilis* endospores also serve as model organisms for the study of germination and sporulation. In these cases, highly purified spores enable more accurate determination of their true protein complement free from confounders from other entities. However, quantitative evidence concerning the purity and proteome composition of *B. subtilis* spores following different harvesting methodologies has been insufficiently reported to date. We observed that although cold water washing only eliminates approximately 8% of vegetative cells and immature spores, it already significantly affected the final proteomic results compared to direct harvesting of spores. Moreover, the most significant distinction was observed between HistodenZ purified spores and directly harvested samples, identifying 280 unique proteins that mainly include membrane proteins and those involved in stress response and exponential and early post-exponential lifestyles. It is reasonable to speculate that proteins from vegetative cells and lysed mother cells may account for a large portion of these proteins, which are not present in mature spores. Analyzing spore samples of differing purities allows for facile determination of proteins that are more likely to stem from ‘contamination’ of vegetative cells in the sample. When observing the relative abundance of proteins in vegetative cells and spores linked to iBAQ rank in vegetative cells (**Figure. 6B**) spore predominant proteins tend to have lower iBAQ ranks then cell predominant proteins in the vegetative cell proteome. Vegetative cell contributions to non-purified or partially purified spores’ samples are usually minor (in the order of 25% DH down to 5% after HistodenZ purification) and protein content between cells and spores does not differ to a significant degree. This means that proteins stemming from residual vegetative cells detected are also more likely to stem from the abundant part of the cellular protein population. This means these proteins have a high iBAQ rank in vegetative cells, conversely in the spore sample they would likely have a low rank. This is an additional way to distinguish whether quantified proteins are likely to stem from residual vegetative cells. This can be of interest in organisms where obtaining high purity spore samples is difficult or impossible. Collectively, the washing step and HistodenZ purification procedures ensure the harvesting of high-purity spores for subsequent proteomic studies, with the washed spores (WS) as favorite for time-efficient dynamic proteomic spore analyses.

Spores from *Bacillus* species are recognized for their resistance to diverse environmental challenges. Its life cycle is composed of three distinct physiological stages with different characteristics. To better elucidate their varied compositions and the representation of shared proteins that constitute the fundamental ’survival kit’ for life, an in-depth proteomic analysis was conducted. By integrating HpH HPLC fractionation methodology with SPEED sample processing, we observed that 63.3% of the quantified proteins were common to both spores and cells, which is a similar finding as in our previous work using metabolic ^15^N labeling for relative quantification. In our prior study we identified only half the number of proteins compared to our current work, underscoring both the advances in mass spectrometry technology as well as the benefits of using non-mechanical disruption and label free quantitation (13). Within the commonly identified proteins, we observed that 49% of the quantified proteins exhibit consistent abundance levels in both vegetative cells and spores, which is also supported by prior data (13). Moreover, by looking at absolute abundance in spores it is clear that core metabolic enzymes, the transcriptional and translational machinery make up abundant parts of the spore proteome, far exceeding what could be explained by cellular ‘contamination’. This corroborates the notion of an evolutionary benefit in employing a foundational set of proteins as ‘minimal survival kit’ in the spore stage ready for use when vegetative life resumes.

The shared proteins primarily fall into the category of cytoplasm related proteins, the biophysical properties (including pH maintenance and regulation of osmotic pressure) of which serve as principal determinants of vital cellular processes and adaptive responses. Furthermore, we identified 575 and 259 unique proteins in vegetative cells and spores respectively, underscoring the distinct developmental biology operative in the two distinct phases of the *B. subtilis* life cycle. The predominant and specific proteins found in spores are mainly related to sporulation and hydrolyses, which aligns with their critical roles in resistance to harsh conditions and use of macromolecules through germination. The sporulation proteins are primarily involved in the formation and maturation of bacterial spores. These proteins ensure the proper packaging and condensation of DNA, as well as the assembly of the highly resistant outer layers that protect the spore’s genetic material (11). On the other hand, the hydrolase-related proteins identified in spores serve a critical function during the germination process - the transition from the dormant spore state back to a metabolically active vegetative cell (52). The predominant proteins were further examined by KEGG pathway analysis which showed proteins involved in peptidoglycan (PG) biosynthesis, glycolysis, carbon metabolism and biosynthesis of secondary metabolites are shared between cells and spores albeit that the relative quantities of the proteins vary in both. Both the spores and cells of *B. subtilis* are encased in a robust PG layer, offering structural stability and a shield against environmental adversities (52, 53). During sporulation, a layer of PG cortex is formed, crucial for spore dehydration and reinforcing spore resistance. However, germination demands the degradation of the cortex PG, thereby facilitating the expansion and growth of the bacteria and enabling the resumption of metabolic activity. PbpF and DacF proteins were found to be predominant in spores (**Supplementary Table 5**), both of which are involved in the final stage of remodeling and modification of PG impacting the cross-linking and thereby influencing spore structure and properties (54, 55). Recent findings have highlighted a close coordination between central carbon metabolism and PG cell wall synthesis, which indicates the degree of gluconeogenic and glycolytic flux are likely to have a profound impact on cell wall synthesis (56, 57). The connection between cell wall synthesis and cellular physiology is not unexpected as the former represents a substantial carbon drain in the central metabolic pathway. This is particularly notable in Gram-positive bacteria wherein the cell wall constitutes 10-20% of the total cell mass.

We found that there are 41 and 22 proteins commonly existing in both *B. subtilis* spores and cells of carbon metabolism (such as GapB, YqkJ, MalS and Icd) and glycolysis pathways (such as AldY, Pgi, GalM and AcoC), respectively. Despite the extensive understanding of *B. subtilis* sporulation, numerous intriguing questions remain to be answered. These include understanding the precise roles and timings of synthesis and degradation of the shared proteins and physiological state of the identified pathways in these metabolically inactive spores. Unraveling these mysteries will illuminate how these elements serve as critical ’switches’, efficiently facilitating the transition from the vegetative state to sporulation, and subsequently initiating the rapid ’awakening’ of the spore in response to favorable environmental conditions. The label-free quantitation employed in this study offers robust data sets, attributable to its capacity to encompass an extensive range of protein abundance. This becomes particularly beneficial when identifying proteins of both minimal and maximal abundance levels and eliminates reliance on metabolic labeling efficiency. Notably, while label-free methodologies necessitate rigorous reproducibility in both sample preparation and mass spectrometry, we have shown the consistent SPEED sample preparation combined with the sensitive and robust TIMS-TOF mass spectrometry ensures such needs are met.

## Conclusion

Overall, our study compared the utility of the OP, SP3 and SPEED sample preparation for *B. subtilis* proteomics. The findings also suggested that the limitation in samples size for the SPEED procedure that still allows desirable protein identification and high repeatability data can be lowered to 1 OD unit. Additionally, the employment of HistodenZ purification for spore samples can achieve high-fidelity spore proteome profiles. Analysis of differentially pure spore samples as well as using iBAQ ranks of vegetative cell proteomes can pinpoint proteins that are more likely to stem from residual vegetative cells in mixed samples. These insights pave the way for the future advancements in bacterial and spore proteomics. The analysis also outlines a proteomic adaptation of *B. subtilis* when they transfer life forms during different environmental conditions. Proteins present in both cells and spores comprise a basic protein set that is essential for life continuity in dormant spores. These proteins participate in key processes including peptidoglycan biosynthesis, glycolysis, carbon metabolism, and secondary metabolite biosynthesis.

## Funding

This work was supported by a PhD studentship (201906170092) from the Chinese Scholarship Council for Y.H.

## Author contributions

Conceptualization, G.K.; methodology, Y.H., W.R., A.D.K, Z.T. and X.G.; formal analysis, Y.H.; writing—original draft preparation, Y.H.; writing—review and editing, S.B., P.S., and G.K. All authors have read and agreed to the published version of the manuscript.

## Declaration of competing interest

The authors declare no conflict of interest.

## Data Availability

All mass spectral data have been deposited into Massive (https://massive.ucsd.edu/ProteoSAFe/static/massive.jsp), with the identifier MSV000093602.

## Notes

### Competing Interest Statement

The authors have declared no competing interest.

### Summary of Updates

Revision of a number of small typo's and grammar errors in the text which were missed. Addition of a missing current address affiliation to one of the authors.

## References

1. Errington J. 2003. Regulation of endospore formation in Bacillus subtilis. Nat Rev Microbiol 1:117–126.

2. Driks A. 2002. Overview: Development in bacteria: spore formation in Bacillus subtilis. Cell Mol Life Sci CMLS 59:389–391.

3. Kovács ÁT. 2019. Bacillus subtilis. Trends Microbiol 27:724–725.

4. Gao X, Swarge BN, Roseboom W, Wang Y, Dekker HL, Setlow P, Brul S, Kramer G. 2022. Changes in the Spore Proteome of Bacillus cereus in Response to Introduction of Plasmids. Microorganisms 10:1695.

5. Tu Z, Dekker HL, Roseboom W, Swarge BN, Setlow P, Brul S, Kramer G. 2021. High Resolution Analysis of Proteome Dynamics during Bacillus subtilis Sporulation. Int J Mol Sci 22:9345.

6. Abhyankar WR, Kamphorst K, Swarge BN, van Veen H, van der Wel NN, Brul S, de Koster CG, de Koning LJ. 2016. The Influence of Sporulation Conditions on the Spore Coat Protein Composition of Bacillus subtilis Spores. Front Microbiol 7.

7. Ravikumar V, Nalpas NC, Anselm V, Krug K, Lenuzzi M, Šestak MS, Domazet-Lošo T, Mijakovic I, Macek B. 2018. In-depth analysis of Bacillus subtilis proteome identifies new ORFs and traces the evolutionary history of modified proteins. Sci Rep 8:17246.

8. Errington J, van der Aart LT. 2020. Microbe Profile: Bacillus subtilis: model organism for cellular development, and industrial workhorse. Microbiology 166:425–427.

9. Abhyankar W, Pandey R, Ter Beek A, Brul S, de Koning LJ, de Koster CG. 2015. Reinforcement of Bacillus subtilis spores by cross-linking of outer coat proteins during maturation. Food Microbiol 45:54–62.

10. Nerber HN, Sorg JA. 2021. The small acid-soluble proteins of Clostridioides difficile are important for UV resistance and serve as a check point for sporulation. PLOS Pathog 17:e1009516.

11. McKenney PT, Driks A, Eichenberger P. 2013. The Bacillus subtilis endospore: assembly and functions of the multilayered coat. 1. Nat Rev Microbiol 11:33–44.

12. Gao X, Swarge BN, Roseboom W, Setlow P, Brul S, Kramer G. 2022. Time-Resolved Proteomics of Germinating Spores of Bacillus cereus. Int J Mol Sci 23:13614.

13. Swarge B, Abhyankar W, Jonker M, Hoefsloot H, Kramer G, Setlow P, Brul S, de Koning LJ. 2020. Integrative Analysis of Proteome and Transcriptome Dynamics during Bacillus subtilis Spore Revival. mSphere 5:e00463–20.

14. Zheng L, Abhyankar W, Ouwerling N, Dekker HL, van Veen H, van der Wel NN, Roseboom W, de Koning LJ, Brul S, de Koster CG. 2016. Bacillus subtilis Spore Inner Membrane Proteome. J Proteome Res 15:585–594.

15. Stelder SK, Mahmud SA, Dekker HL, de Koning LJ, Brul S, de Koster CG. 2015. Temperature Dependence of the Proteome Profile of the Psychrotolerant Pathogenic Food Spoiler Bacillus weihenstephanensis Type Strain WSBC 10204. J Proteome Res 14:2169–2176.

16. Liu H, Ray WK, Helm RF, Popham DL, Melville SB. 2016. Analysis of the Spore Membrane Proteome in Clostridium perfringens Implicates Cyanophycin in Spore Assembly. J Bacteriol 198:1773–1782.

17. Li L, Jin J, Hu H, Deveau IF, Foley SL, Chen H. 2022. Optimization of sporulation and purification methods for sporicidal efficacy assessment on Bacillus spores. J Ind Microbiol Biotechnol 49:kuac014.

18. de Bruin OM, Birnboim HC. 2016. A method for assessing efficiency of bacterial cell disruption and DNA release. BMC Microbiol 16:197.

19. Swarge BN, Roseboom W, Zheng L, Abhyankar WR, Brul S, de Koster CG, de Koning LJ. 2018. “One-Pot” Sample Processing Method for Proteome-Wide Analysis of Microbial Cells and Spores. Proteomics Clin Appl 12:e1700169.

20. Protein Analysis by Shotgun/Bottom-up Proteomics | Chemical Reviews. https://pubs.acs.org/doi/full/10.1021/cr3003533. Retrieved 10 July 2023.

21. Leitner A, Aebersold R. 2013. SnapShot: mass spectrometry for protein and proteome analyses. Cell 154:252–252.e1.

22. Kramer G, Woolerton Y, van Straalen JP, Vissers JP, Dekker N, Langridge JI, Beynon RJ, Speijer D, Sturk A, Aerts JM. 2015. Accuracy and reproducibility in quantification of plasma protein concentrations by mass spectrometry without the use of isotopic standards. PLoS One 10:e0140097.

23. Kulak NA, Pichler G, Paron I, Nagaraj N, Mann M. 2014. Minimal, encapsulated proteomic-sample processing applied to copy-number estimation in eukaryotic cells. Nat Methods 11:319–324.

24. Wiśniewski JR, Zougman A, Nagaraj N, Mann M. 2009. Universal sample preparation method for proteome analysis. 5. Nat Methods 6:359–362.

25. Hughes CS, Foehr S, Garfield DA, Furlong EE, Steinmetz LM, Krijgsveld J. 2014. Ultrasensitive proteome analysis using paramagnetic bead technology. Mol Syst Biol 10:757.

26. Petruschke H, Anders J, Stadler PF, Jehmlich N, von Bergen M. 2020. Enrichment and identification of small proteins in a simplified human gut microbiome. J Proteomics 213:103604.

27. Sielaff M, Kuharev J, Bohn T, Hahlbrock J, Bopp T, Tenzer S, Distler U. 2017. Evaluation of FASP, SP3, and iST Protocols for Proteomic Sample Preparation in the Low Microgram Range. J Proteome Res 16:4060–4072.

28. Wang C-Y, Lempp M, Farke N, Donati S, Glatter T, Link H. 2021. Metabolome and proteome analyses reveal transcriptional misregulation in glycolysis of engineered E. coli. 1. Nat Commun 12:4929.

29. Huang Y, Swarge BN, Roseboom W, Bleeker JD, Brul S, Setlow P, Kramer G. 2023. Integrative metabolomics and proteomics allow the global intracellular characterization of Bacillus subtilis cells and spores. bioRxiv 10.1101/2023.06.19.545067.

30. Doellinger J, Schneider A, Hoeller M, Lasch P. 2020. Sample Preparation by Easy Extraction and Digestion (SPEED) - A Universal, Rapid, and Detergent-free Protocol for Proteomics Based on Acid Extraction. Mol Cell Proteomics 19:209–222.

31. Abele M, Doll E, Bayer FP, Meng C, Lomp N, Neuhaus K, Scherer S, Kuster B, Ludwig C. 2023. Unified Workflow for the Rapid and In-Depth Characterization of Bacterial Proteomes. Mol Cell Proteomics 22.

32. Kort R, O’Brien AC, van Stokkum IHM, Oomes SJCM, Crielaard W, Hellingwerf KJ, Brul S. 2005. Assessment of Heat Resistance of Bacterial Spores from Food Product Isolates by Fluorescence Monitoring of Dipicolinic Acid Release. Appl Environ Microbiol 71:3556–3564.

33. Ghosh S, Korza G, Maciejewski M, Setlow P. 2015. Analysis of Metabolism in Dormant Spores of Bacillus Species by 31P Nuclear Magnetic Resonance Analysis of Low-Molecular-Weight Compounds. J Bacteriol 197:992–1001.

34. Wu T, Hu E, Xu S, Chen M, Guo P, Dai Z, Feng T, Zhou L, Tang W, Zhan L, Fu X, Liu S, Bo X, Yu G. 2021. clusterProfiler 4.0: A universal enrichment tool for interpreting omics data. The Innovation 2:100141.

35. Nagaraj N, Wisniewski JR, Geiger T, Cox J, Kircher M, Kelso J, Pääbo S, Mann M. 2011. Deep proteome and transcriptome mapping of a human cancer cell line. Mol Syst Biol 7:548.

36. Zeiler M, Moser M, Mann M. 2014. Copy number analysis of the murine platelet proteome spanning the complete abundance range. Mol Cell Proteomics 13:3435–3445.

37. Wiśniewski JR, Hein MY, Cox J, Mann M. 2014. A “proteomic ruler” for protein copy number and concentration estimation without spike-in standards. Mol Cell Proteomics 13:3497–3506.

38. Silva JC, Gorenstein MV, Li G-Z, Vissers JP, Geromanos SJ. 2006. Absolute quantification of proteins by LCMSE: A virtue of parallel MS acquisition* S. Mol Cell Proteomics 5:144–156.

39. Wiśniewski JR, Rakus D. 2014. Multi-enzyme digestion FASP and the ‘Total Protein Approach’-based absolute quantification of the Escherichia coli proteome. J Proteomics 109:322–331.

40. Schwanhäusser B, Busse D, Li N, Dittmar G, Schuchhardt J, Wolf J, Chen W, Selbach M. 2011. Global quantification of mammalian gene expression control. Nature 473:337–342.

41. Armengaud J. 2016. Next-generation proteomics faces new challenges in environmental biotechnology. Curr Opin Biotechnol 38:174–182.

42. Armengaud J. 2013. Microbiology and proteomics, getting the best of both worlds! Environ Microbiol 15:12–23.

43. Miller RM, Smith LM. 2023. Overview and considerations in bottom-up proteomics. Analyst 148:475–486.

44. de Boer R, Peters R, Gierveld S, Schuurman T, Kooistra-Smid M, Savelkoul P. 2010. Improved detection of microbial DNA after bead-beating before DNA isolation. J Microbiol Methods 80:209–211.

45. Varnavides G, Madern M, Anrather D, Hartl N, Reiter W, Hartl M. 2022. In Search of a Universal Method: A Comparative Survey of Bottom-Up Proteomics Sample Preparation Methods. J Proteome Res 21:2397–2411.

46. Templeton EM, Pilbrow AP, Kleffmann T, Pickering JW, Rademaker MT, Scott NJA, Ellmers LJ, Charles CJ, Endre ZH, Richards AM, Cameron VA, Lassé M. 2023. Comparison of SPEED, S-Trap, and In-Solution-Based Sample Preparation Methods for Mass Spectrometry in Kidney Tissue and Plasma. 7. Int J Mol Sci 24:6290.

47. Zhang X. 2015. Less is More: Membrane Protein Digestion Beyond Urea–Trypsin Solution for Next-level Proteomics*. Mol Cell Proteomics 14:2441–2453.

48. Wood TK, Knabel SJ, Kwan BW. 2013. Bacterial Persister Cell Formation and Dormancy. Appl Environ Microbiol 79:7116–7121.

49. Boxberger M, Cenizo V, Cassir N, La Scola B. 2021. Challenges in exploring and manipulating the human skin microbiome. Microbiome 9:125.

50. Pierce CG, Uppuluri P, Tristan AR, Wormley FL, Mowat E, Ramage G, Lopez-Ribot JL. 2008. A simple and reproducible 96 well plate-based method for the formation of fungal biofilms and its application to antifungal susceptibility testing. Nat Protoc 3:1494–1500.

51. Piehowski PD, Petyuk VA, Orton DJ, Xie F, Moore RJ, Ramirez-Restrepo M, Engel A, Lieberman AP, Albin RL, Camp DG, Smith RD, Myers AJ. 2013. Sources of technical variability in quantitative LC-MS proteomics: human brain tissue sample analysis. J Proteome Res 12:2128–2137.

52. Brogan AP, Rudner DZ. 2023. Regulation of peptidoglycan hydrolases: localization, abundance, and activity. Curr Opin Microbiol 72:102279.

53. Rohs PDA, Bernhardt TG. 2021. Growth and Division of the Peptidoglycan Matrix. Annu Rev Microbiol 75:315–336.

54. Popham DL, Bernhards CB. 2015. Spore Peptidoglycan. Microbiol Spectr 3:10.1128/microbiolspec.tbs-0005-2012.

55. Popham DL, Gilmore ME, Setlow P. 1999. Roles of Low-Molecular-Weight Penicillin-Binding Proteins in Bacillus subtilis Spore Peptidoglycan Synthesis and Spore Properties. J Bacteriol 181:126–132.

56. Kawai Y, Mercier R, Mickiewicz K, Serafini A, Sório de Carvalho LP, Errington J. 2019. Crucial role for central carbon metabolism in the bacterial L-form switch and killing by β-lactam antibiotics. Nat Microbiol 4:1716–1726.

57. Sperber AM, Herman JK. 2017. Metabolism Shapes the Cell. J Bacteriol 199:e00039–17.

