## Supplementary figures for "Non-mechanical disruption of *Bacillus subtilis spores* allows sensitive and deep characterization of the minimal proteome for resuming a cellular lifestyle"

**Supplemental Figures**

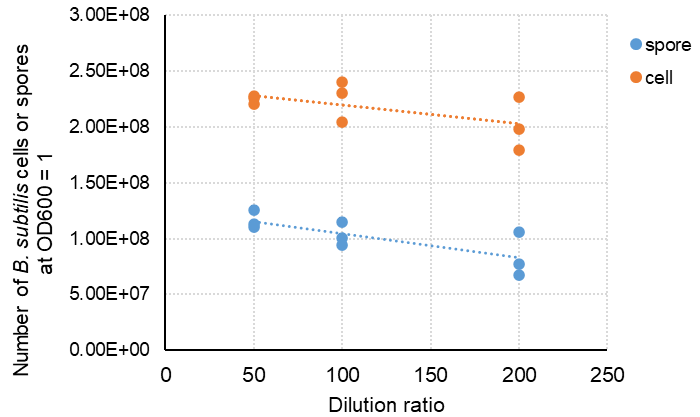

**Figure S1. Numbers of *B. subtilis* cells or spores in solutions with an OD_600_ of 1 at different dilution ratios.**

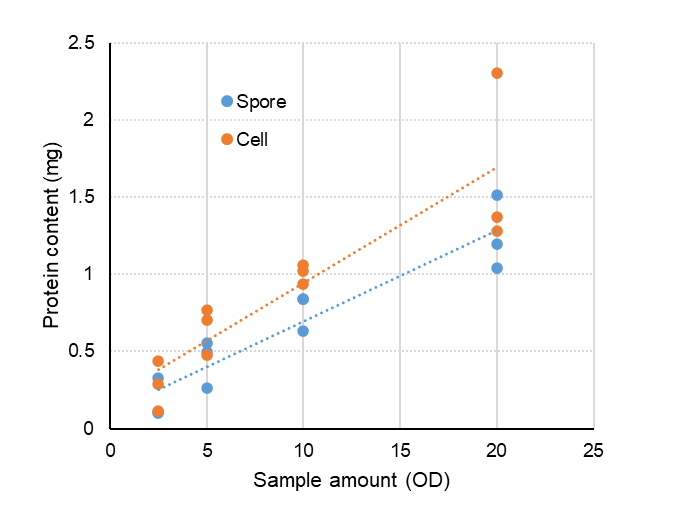

**Figure S2. Relation between Sample amount in OD600 and the protein content in mg determined by BCA assay (assuming 1 ml of solution).**

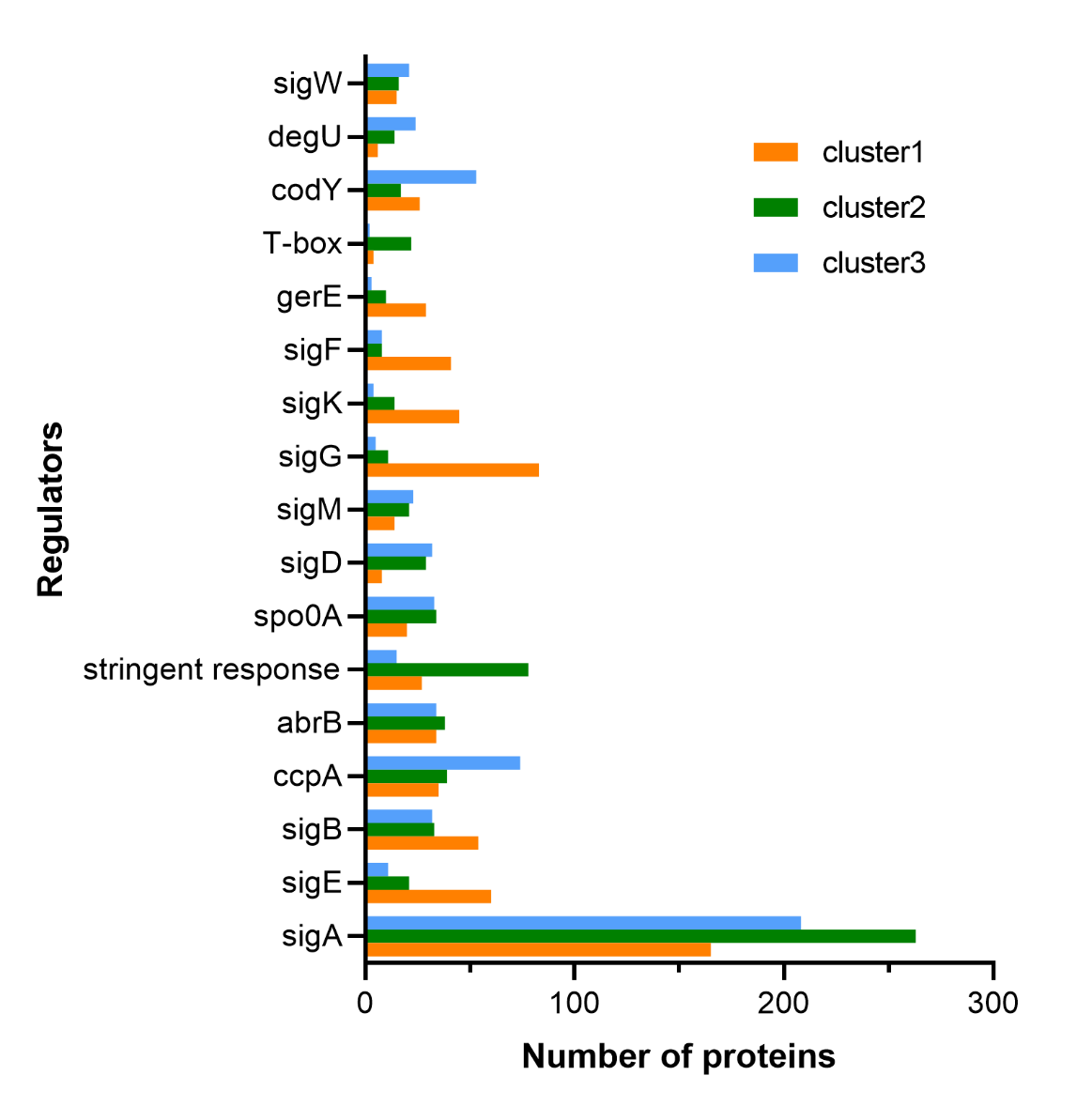

**Figure S3. Distribution of protein counts among the top 10 regulators in each cluster from figure 5 in the main text.**

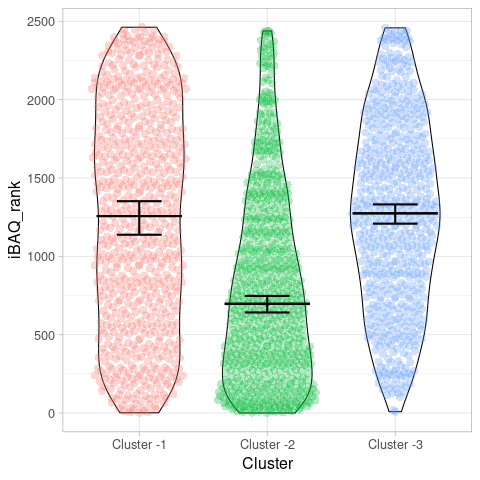

**Figure S4. Violin plot of iBAQ rank in vegetative cells for proteins found in the clusters shown in Figure 5 in the main text.**
